# Transiently boosting Vγ9+Vδ2+ γδ T cells early in Mtb coinfection of SIV-infected juvenile macaques does not improve Mtb host resistance

**DOI:** 10.1101/2024.07.22.604654

**Authors:** Erica C. Larson, Amy L. Ellis, Mark A. Rodgers, Abigail K. Gubernat, Janelle L. Gleim, Ryan V. Moriarty, Alexis J. Balgeman, Yonne T. de Menezes, Cassaundra L. Ameel, Daniel J. Fillmore, Skyler M. Pergalske, Jennifer A. Juno, Pauline Maiello, Harris B. Chishti, Philana Ling Lin, Dale I. Godfrey, Stephen J. Kent, Daniel G. Pellicci, Lishomwa C. Ndhlovu, Shelby L. O’Connor, Charles A. Scanga

**Affiliations:** Department of Microbiology and Molecular Genetics, University of Pittsburgh School of Medicine, Pittsburgh, Pennsylvania, USA; Center for Vaccine Research, University of Pittsburgh School of Medicine, Pittsburgh, Pennsylvania, USA; Department of Pathology and Laboratory Medicine, University of Wisconsin - Madison, Wisconsin, USA; Department of Immunobiology, Federal University of Santa Catarina, Florianópolis, Santa Catarina, Brazil; Department of Microbiology and Immunology, The Peter Doherty Institute for Infection and Immunity, University of Melbourne, Melbourne, VIC, Australia; Department of Pediatrics, UPMC’s Children’s Hospital of the University of Pittsburgh of UPMC, Pittsburgh, PA; Melbourne Sexual Health Centre and Department of Infectious Diseases, Alfred Hospital and Centre Clinical School, Monash University, Melbourne, VIC, Australia; Department of Paediatrics, University of Melbourne, Melbourne, VIC, Australia; Department of Medicine, Division of Infectious Disease, Weill Cornell Medicine, New York, New York, USA; Wisconsin National Primate Research Center, University of Wisconsin - Madison, Wisconsin, USA

## Abstract

Children living with HIV have a higher risk of developing tuberculosis (TB), a disease caused by the bacterium *Mycobacterium tuberculosis* (Mtb). Gamma delta (γδ) T cells in the context of HIV/Mtb coinfection have been understudied in children, despite *in vitro* evidence suggesting γδ T cells assist with Mtb control. We investigated whether boosting a specific subset of γδ T cells, phosphoantigen-reactive Vγ9+Vδ2+ cells, could improve TB outcome using a nonhuman primate model of pediatric HIV/Mtb coinfection. Juvenile Mauritian cynomolgus macaques (MCM), equivalent to 4–8-year-old children, were infected intravenously (i.v.) with SIV. After 6 months, MCM were coinfected with a low dose of Mtb and then randomized to receive zoledronate (ZOL), a drug that increases phosphoantigen levels, (n=5; i.v.) at 3- and 17-days after Mtb accompanied by recombinant human IL-2 (s.c.) for 5 days following each ZOL injection. A similarly coinfected MCM group (n=5) was injected with saline as a control. Vγ9+Vδ2+ γδ T cell frequencies spiked in the blood, but not airways, of ZOL+IL-2-treated MCM following the first dose, however, were refractory to the second dose. At necropsy eight weeks after Mtb, ZOL+IL-2 treatment did not reduce pathology or bacterial burden. γδ T cell subset frequencies in granulomas did not differ between treatment groups. These data show that transiently boosting peripheral γδ T cells with ZOL+IL-2 soon after Mtb coinfection of SIV-infected MCM did not improve Mtb host defense.

## Introduction

Children living with HIV have a higher risk of developing tuberculosis (TB) even when virally suppressed with antiretroviral therapy (ART) (1–5). TB disease is often more severe in children (*e.g.*, miliary TB and TB meningitis) (6–8) and TB-associated mortality is higher in children living with HIV than those without HIV (9). Although TB is curable, the consequence of disease on lung function after the bacteria have been cleared is a growing concern, especially in children (10). TB early in life could have long-term health consequences due to the pulmonary and extrapulmonary damage sustained during infection. However, very few therapies exist to prevent or reverse this tissue damage. Host-directed therapy, which modulates the host immune response to eradicate Mtb, is a promising strategy that may minimize TB disease and subsequent tissue damage in children, especially those living with HIV.

Gamma delta (γδ) T cells are a subset of unconventional T cells that typically recognize non-peptide antigens (11). In addition to direct effector functions, γδ T cells crosstalk with several immune cell types and provide a variety of supportive functions including B cell interactions, maturation of dendritic cells, activation of neutrophils as well as NK cells, and priming of conventional αβ T cells (12). One subtype of γδ T cells, Vγ9+Vδ2+ γδ T cells, which react against metabolites from the isoprenoid synthesis pathway known as phosphoantigens (13), display both direct killing of bacilli and indirect antimicrobial activity by enhancing CD4+ and CD8+ T cell responses against Mtb both *in vitro* and *in vivo* (14–16). Augmenting Vγ9+Vδ2+ γδ T cells has been shown to reduce Mtb burden in rhesus macaques (16–18). In humans, Vδ2+ γδ T cells frequencies are highly dynamic throughout childhood (19–21). Vδ2+ γδ T cells peak in blood during pre-adolescence (5-9 years old) then slowly decline into adulthood (19). It is not well understood how these changes in γδ T cell frequencies throughout development influence disease control of pathogens like HIV and Mtb in children. Circulating Vδ2+ γδ T cells are notably depleted in people living with HIV, perhaps due to their higher expression of CCR5 (11, 22–24). Evidence suggests that the Vδ2+ γδ T cells that do remain in individuals living with HIV are anergic, especially to mycobacterial antigens (25). It has yet to be determined whether HIV-mediated Vδ2+ γδ T cell depletion contributes to increased TB susceptibility and whether pharmacologic modulation of this cell population would reduce TB disease in children.

Using our juvenile nonhuman primate (NHP) model of pediatric HIV/Mtb coinfection (26), we evaluated whether boosting Vγ9+Vδ2+ γδ T cells reduces TB disease. Zoledronate (ZOL; brand name, Reclast®), is an FDA-approved drug used for treating osteoporosis, Paget’s disease of bone, and high levels of calcium in the blood caused by certain types of cancer (27). ZOL is known to increase cellular phosphoantigen levels by blocking the isoprenoid biosynthesis pathway (13). In combination with IL-2, ZOL can increase Vγ9+Vδ2+ γδ T cell frequency *in vitro* and *in vivo* (18, 28–30). We administered ZOL+IL-2 to SIV-infected juvenile macaques at 3 and 17 days after Mtb coinfection. Vγ9+Vδ2+ γδ T cells transiently increased in the blood following the first dose of ZOL+IL-2 but were refractory to the second dose. ZOL+IL-2 did not increase γδ T cell frequencies in airways or in TB granulomas. TB outcome did not improve with ZOL+IL-2 treatment in SIV/Mtb coinfected juvenile macaques, suggesting that transient changes to Vγ9+Vδ2+ γδ T cells in the periphery early in Mtb coinfection did not improve host immunity to Mtb.

## Results

### ZOL+IL-2 does not alter plasma SIV viremia

Ten juvenile macaques (aged 1-2 years old, equivalent to 4–8 year-old human children) were intravenously infected with SIVmac239M (Figure 1A). After six months, animals were infected with a low dose of virulent Mtb via bronchoscope. A subset of animals (n = 5) was given zoledronate (ZOL; 0.2. mg/kg, i.v.) at days 3 and 17 after Mtb coinfection. ZOL-treated animals also received recombinant human IL-2 subcutaneously once daily (SID) for 5 days after each ZOL administration. The remaining five animals served as a control group and were given saline.

**Figure 1.**
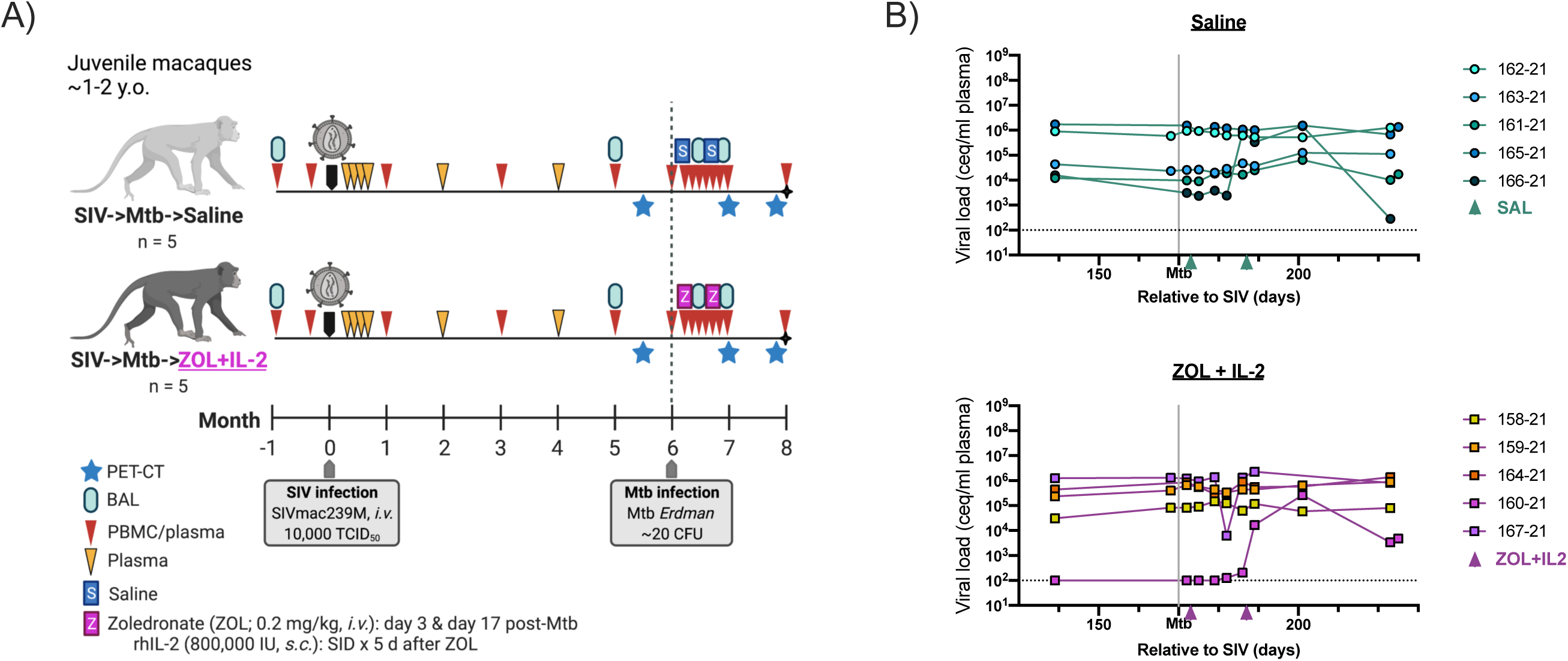
Study timeline and plasma viremia. A) Study timeline. B) Plasma viral load (viral copy equivalents/mL) were determined by qRT-PCR. Each point indicates an individual animal. Triangles along x-axis indicate when treatment was administered (days 3 and day 17 post Mtb). Horizontal dashed line represents the limit of detection (100 ceq/mL plasma). Mixed effects models (two-tailed) with subject as a random variable were used to assess mean differences among time points and treatment groups. No significant differences were determined between time or treatment (p = 0.4979 and p = 0.9711, respectively).

Plasma SIV viremia was measured throughout the entire study. All 10 animals exhibited peak viremia ∼2 weeks post SIV infection followed by stabilization at setpoints unique to each animal (Figure S1). One animal (160-21), randomized to the ZOL+IL-2 group, was a spontaneous viral controller with viremia that eventually dropped below the limit of detection (100 ceq/mL). Both saline control and ZOL+IL-2 groups had stable plasma viral setpoints prior to Mtb coinfection (Figure 1B). Following Mtb coinfection and ZOL+IL-2 or saline administration, plasma viral load did not significantly differ between the two groups. Plasma viremia was transiently elevated after Mtb coinfection in just two animals: 166-21 (saline) and 160-21 (ZOL+IL-2). This burst of SIV replication was not observed in most animals and, thus, did not appear to be related to ZOL+IL-2 treatment.

### ZOL+IL-2 induces a transient spike of circulating Vγ9+Vδ2+ γδ T cells

The frequency of circulating Vγ9+Vδ2+ γδ T cells transiently increased in ZOL+IL-2-treated animals shortly after the first dose, between days 7 and 10 after Mtb coinfection (Figure 2A). Peak Vγ9+Vδ2+ γδ T cell frequencies were significantly higher in ZOL+IL-2-treated animals compared to saline-treated animals (Figure 2B). However, Vγ9+Vδ2+ γδ T cell frequencies returned to their original levels 14 days after Mtb coinfection and did not re-expand in response to the second dose of ZOL+IL-2 at day 17 after Mtb coinfection. Vγ9+Vδ2+ γδ T cells did not expand in the saline control group (Figure 2A & B). In airways, sampled by bronchoalveolar lavage (BAL), ZOL+IL-2 had little effect on Vγ9+Vδ2+ γδ T cell frequencies, although one animal 160-21 showed an increase in Vγ9+Vδ2+ γδ T cell after drug treatment (Figure 2C). Conventional CD4+ and CD8+ T cell subsets did not significantly differ in the blood or airways between the two groups following drug treatment (Figure S2). Gating strategies for circulating and airway T cell subsets are shown in Figure S3 and S4A, respectively.

**Figure 2.**
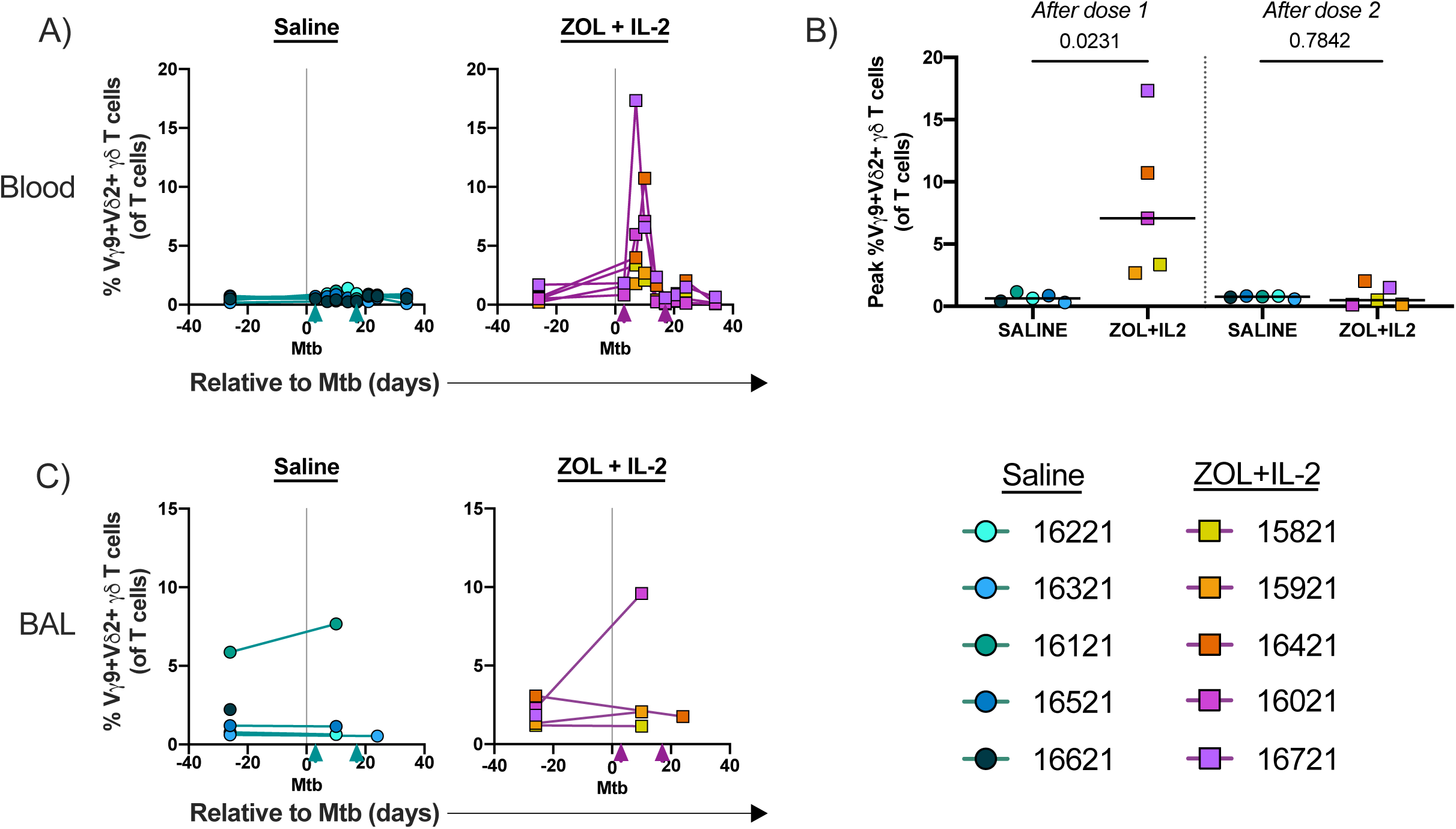
Vγ9+Vδ2+ γδ T cells in blood and BAL after Mtb coinfection and ZOL+IL-2/saline treatment. Individual symbols indicate individual animals. Triangles along x-axis indicate when treatment was administered (days 3 and day 17 post Mtb). A) Frequencies of Vγ9+Vδ2+ γδ T cells in blood from saline-treated (left panel) and ZOL+IL-2-treated (right panel) animals. B) Statistical comparison of peak Vγ9+Vδ2+ γδ T cells following dose 1 (Post Mtb d3) and dose 2 (Post Mtb d17). Unpaired t tests were performed to determine significance. P-values are shown. C) Frequencies of Vγ9+Vδ2+ γδ T cells in BAL from saline-treated (left panel) and ZOL+IL-2-treated (right panel) animals.

### ZOL+IL-2 does not alter T cell composition in granulomas, but may alter cytotoxic profiles

Next, we sought to determine whether the transient spike of circulating Vγ9+Vδ2+ γδ T cells altered the cellular composition of granulomas, the sites of Mtb infection. We characterized γδ T cell subsets as well as CD4+, CD8αβ+, and CD8αα+ T cells in granulomas harvested at necropsy, 8 weeks after Mtb coinfection. γδ T cell subset frequencies did not differ between the two treatment groups (Figure 3A-D). There were also no differences in the CD4+ and CD8+ T cell subset frequencies between the two treatment groups (Figure 3E-I). Due to the low numbers of Vγ9+Vδ2+ γδ T cells within granulomas (<50 events/sample), we were unable to characterize cytokine production and cytotoxic profiles in this population. In contrast, CD8αβ+ and CD8αα+ T cells in granulomas were numerous enough to more fully characterize.

**Figure 3.**
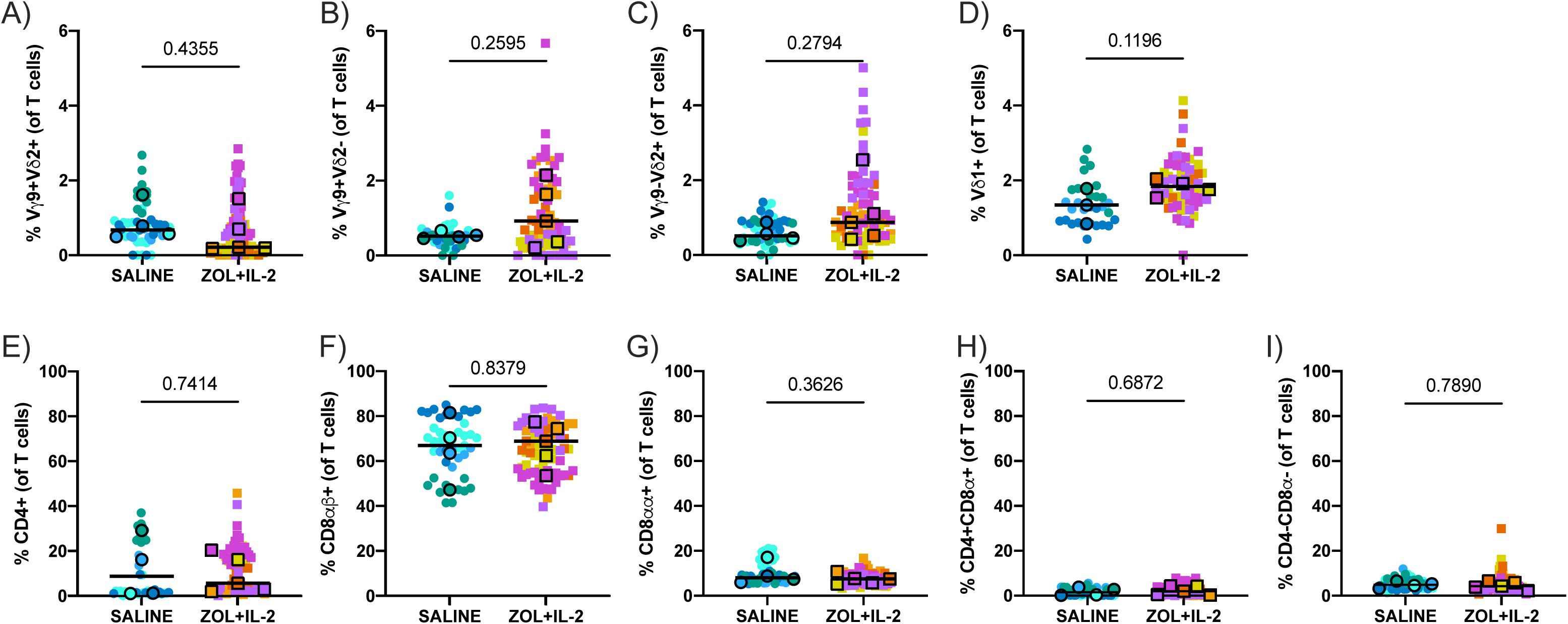
T cell composition in granulomas from saline and ZOL+IL-2-treated animals. Frequencies of T cell subsets in granulomas relative to T cell (CD3+) gate. A) Vγ9+Vδ2+ T cells. B) Vγ9+Vδ2-T cells. C) Vγ9-Vδ2+ T cells. D) Vδ1+ T cells. E) CD4+ T cells. F) CD8ɑβ+ T cells. G) CD8ɑɑ+ T cells. H) CD4+CD8ɑ+ T cells. I) CD4-CD8ɑ-T cells. Outlined symbols indicate median per animal and unlined symbols indicate individual samples. Bars indicate group medians. Two tailed, unpaired t tests of group medians were performed to determine significance. *P*-values are shown.

Frequencies of granulysin (Glyn+)-positive CD8αβ+ T cells, but not granzyme B (GrzB+) or perforin (Perf+), were significantly higher in granulomas from ZOL+IL2-treated animals (Figure 4A-C). Similarly, CD8αα+ T cells in granulomas from ZOL+IL-2-treated animals had higher frequencies of Glyn+, while GrzB+ frequencies were statistically significantly lower (Figure 4D-F). Cytotoxic profiles by Boolean gating of CD8αβ+ and CD8αα+ T cells in granulomas revealed that the profile from ZOL+IL-2-treated animals differed from saline-treated animals (Figure 5A-B). There were significantly higher frequencies of Glyn+, GrzB+, and Perf+ triple-positive as well as Glyn+Perf+ double-positive CD8αβ+ T cells in granulomas from ZOL+IL-2-treated animals, while single-positive Perf+ frequencies were significantly lower (Figure 5C-E). CD8αα+ T cells in granulomas from ZOL+IL-2-treated animals were comprised of significantly higher frequencies of Glyn+Perf+ and Glyn+ populations compared to granulomas from saline-treated controls (Figure 5F-G). We also measured *de novo* production of the cytokines IFNγ, TNF, IL-2, and IL-17 from CD8αβ+ and CD8αα+ T cells in granulomas and found that ZOL+IL-2 did not increase the *de novo* production (Figure S5). The gating strategy for granuloma T cell subsets is shown in Figure S6.

**Figure 4.**
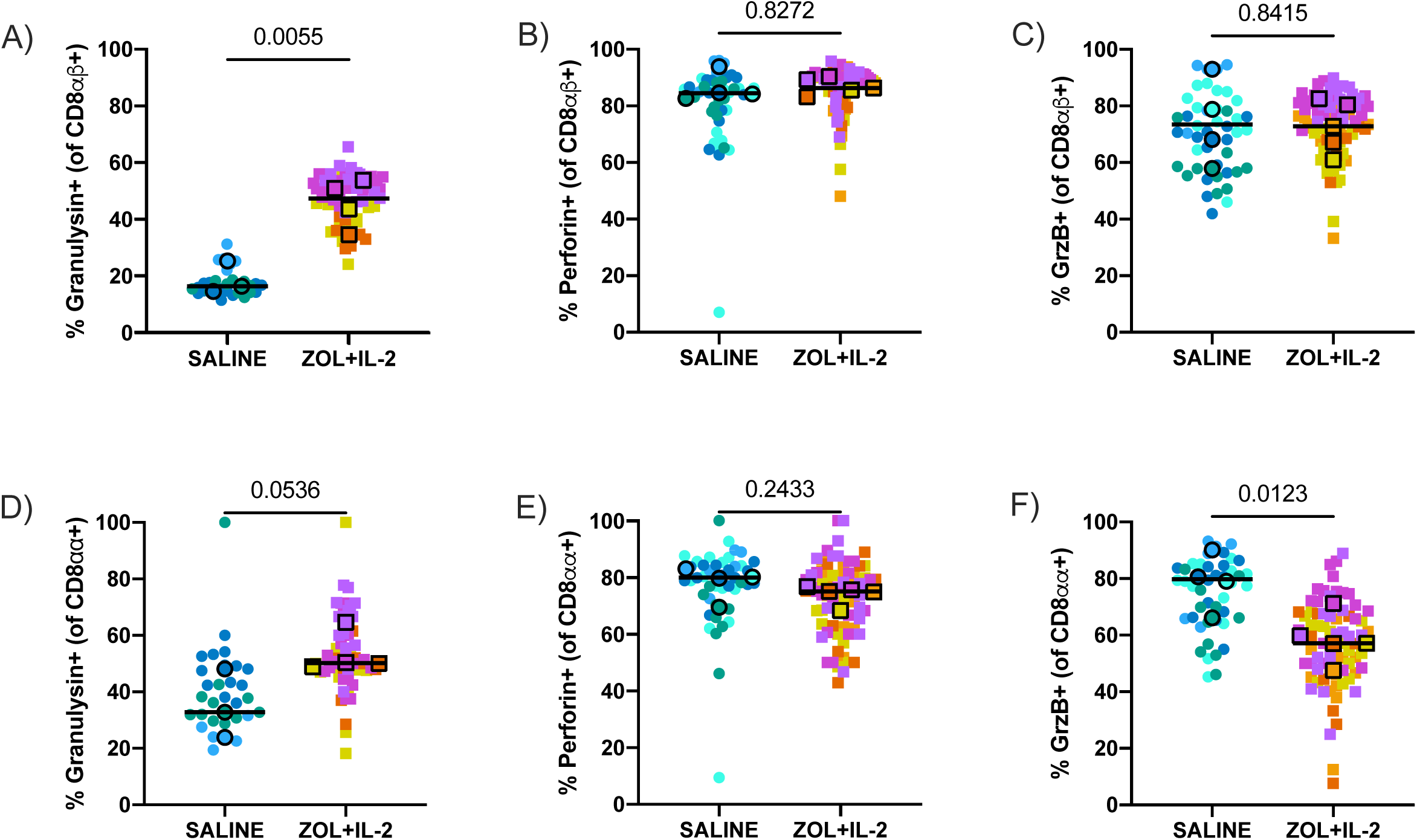
ZOL+IL-2 treatment induces granulysin in CD8ɑβ+ and CD8ɑɑ+ T cells isolated from granulomas but reduces GrzB in CD8ɑɑ+ T cells. A-C) Frequencies of granulysin (A), perforin (B), and GrzB (C) in CD8ɑβ+ T cells. D-F) Frequencies of granulysin (D), perforin (E), and GrzB (F) in CD8ɑɑ+ T cells. Outlined symbols indicate median per animal and unlined symbols indicate individual samples. Bars indicate group medians. Unpaired t tests of group medians were performed to determine significance. P-values are shown.

**Figure 5.**
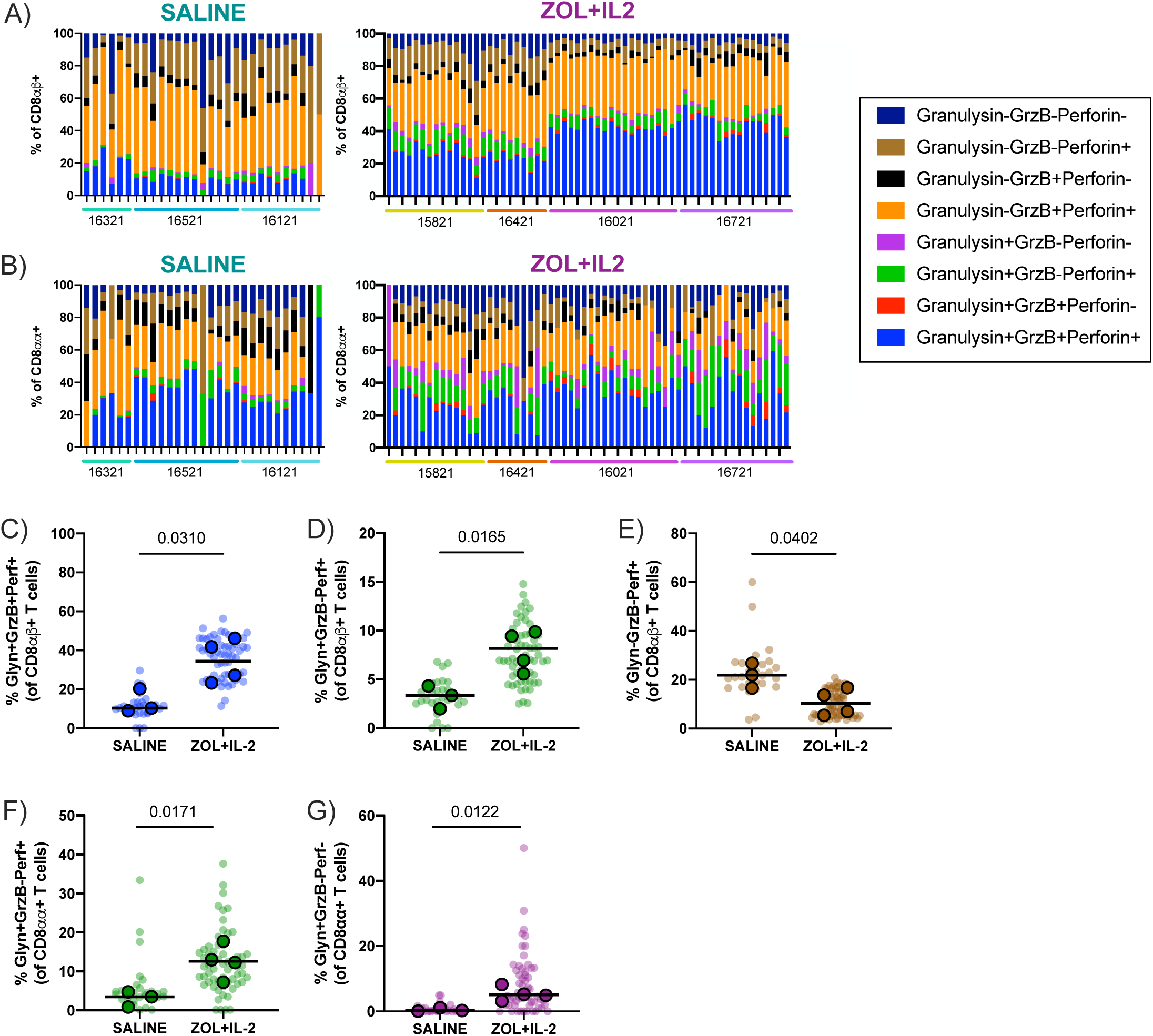
ZOL+IL-2 may alter cytotoxic profiles of granuloma CD8 T cell subsets. A-B) Cytotoxic profiles of CD8ɑβ+ T cells (A) and CD8ɑɑ+ T cells (B) determined by Boolean gating. Each bar indicates an individual granuloma and granulomas from individual animals are indicated on the x-axis. C-E) Frequencies of significant cytotoxic populations of CD8ɑβ+ T cells in granulomas. F-G) Frequencies of significant cytotoxic populations of CD8ɑɑ+ T cells in granulomas. C-G) Outlined symbols indicate median per animal and unlined symbols indicate individual samples. Bars indicate group medians. Unpaired t tests of group medians were performed to determine significance. P-values are shown.

### ZOL+IL-2 does not reduce TB disease

We next assessed whether the transient burst of circulating Vγ9+Vδ2+ γδ T cells after ZOL+IL-2 shortly after Mtb coinfection was associated with improved Mtb control and reduced TB disease. Following Mtb coinfection, we monitored erythrocyte sedimentation rate (ESR), and Mtb culture status of gastric aspirates (GA) and BAL fluid (BALF) (Table S1). Only one animal, assigned to the ZOL+IL-2 group, exhibited a transiently elevated ESR, Mtb was cultured from GA of 3/5 animals in both groups, and BALF was uniformly culture-negative (Table S1). There was no significant change in body weight of any animal (data not shown). Thus, clinical parameters of TB did not differ between the ZOL+IL-2 and saline groups.

Following Mtb coinfection, lung inflammation was measured at 4- and 8-weeks post Mtb by PET/CT imaging. ZOL+IL-2 did not reduce lung inflammation at either time point compared to saline-treated animals (Figure S7). At the time of necropsy around 8-9 weeks after Mtb coinfection (Table S1), we measured lung inflammation, overall TB pathology, and total Mtb burden. These parameters were similar between the ZOL+IL-2- and the saline-treated animals (Figure 6A-C). Likewise, no significant differences were noted between the groups when we assessed lung-specific pathology and Mtb burden (Figure 6D-E). The pathology score for thoracic lymph nodes was significantly higher, due to a higher number of involved lymph nodes, in ZOL+IL-2-treated animals (Figure 6F), although there was no difference in thoracic lymph node Mtb burden (Figure 6G). Extrapulmonary scores did not differ between the two treatment groups (Figure 6H).

**Figure 6.**
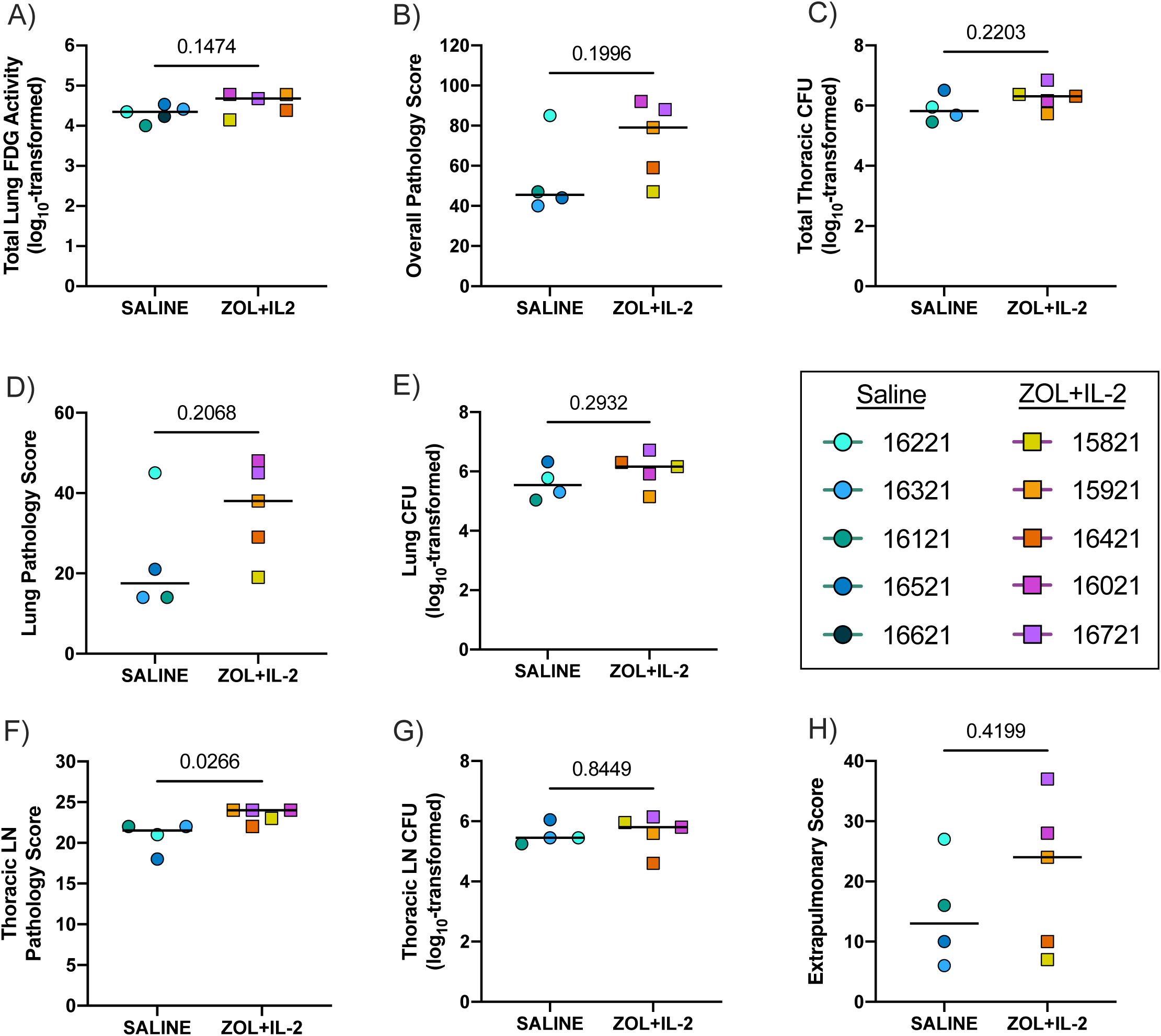
ZOL+IL-2 does not reduce bacterial burden or TB pathology. Symbols indicate individual animals and bars indicate medians of group. Two-tailed, statistical tests of group medians were performed. Unpaired t tests were performed for all. *P*-values are shown. A) Total Lung FDG Activity at necropsy. B) Overall Pathology Score. C) Total Thoracic CFU. D) Lung Pathology Score. E) Lung CFU. F) Thoracic Lymph Nodes (LN) Pathology Score. G) Thoracic LN CFU. H) Extrapulmonary Score.

Overall, these data show that administering ZOL+IL-2 to SIV-infected macaques soon after Mtb coinfection transiently increases the frequency of Vγ9+Vδ2+ γδ T cells in the blood, but not the airway. ZOL+IL-2 did not alter the cellular composition of granulomas, although it did affect the cytotoxic profiles of CD8αβ+ T cells in these lesions. Nonetheless, the effects of ZOL+IL-2 did not significantly alter TB disease or Mtb burden despite these peripheral immunological changes.

## Discussion

In this study, we sought to boost Vγ9+Vδ2+ γδ T cells, a γδ T cell subset associated with anti-Mtb activity (14–16), using our nonhuman primate model of pediatric HIV/Mtb coinfection (26) to determine whether augmenting Vγ9+Vδ2+ γδ T cells could reduce Mtb burden and overall TB disease in the presence of a pre-existing SIV infection. Several groups have shown that inducing Vγ9+Vδ2+ γδ T cells by various approaches is moderately efficacious against Mtb infection in SIV-naïve, adult rhesus macaques (16–18). However, the effect of boosting Vγ9+Vδ2+ γδ T cells against Mtb in the presence of a pre-existing SIV infection, which exacerbates TB disease (31), has yet to be explored. When we administered ZOL+IL-2 three days after Mtb coinfection of MCM chronically infected with SIV, we observed a transient increase of circulating Vγ9+Vδ2+ γδ T cells, but not in the airways. We also observed changes in the cytotoxic profiles of CD8αβ+ and CD8αα+ T cells in granulomas harvested from ZOL+IL-2-treated animals 8 weeks after Mtb coinfection. However, ZOL+IL-2 had little effect on overall Mtb burden or TB progression.

Chen and others have reported transient induction of circulating Vγ9+Vδ2+ γδ T cells occurring 4 to 7 days after initial *in vivo* administration of drugs that increase phosphoantigens, including ZOL (17, 29, 30, 32). Whether transient expansion also occurs within tissue resident Vγ9+Vδ2+ γδ T cell populations was not assessed, although expansion of these cells was reported in airways 8 to 14 days after ZOL treatment (29, 30). In contrast, we did not observe increased numbers of Vγ9+Vδ2+ γδ T cells in the airways following ZOL+IL-2 administration, although it is possible we missed a transient increase if its duration was short. Although we did not measure it here, ZOL+IL-2 treatment may reduce CCR6 expression, an important mediator of immune cell trafficking to the lung (33, 34), on circulating Vγ9+Vδ2+ γδ T cells (30). Decreased CCR6 expression corresponded with increased circulating Vγ9+Vδ2+ γδ T cells (30), suggesting that these circulating cells may have limited ability to traffic to tissues and the expansion observed in airways may be due to proliferation of local tissue resident cell populations. However, in another study (18), transfer of *ex vivo*-expanded autologous Vγ9+Vδ2+ γδ T cells did, in fact, traffic to airways and were retained several weeks later. Further studies of Vγ9+Vδ2+ γδ T cell trafficking and retention within various tissue compartments would provide insights into improving the efficacy of γδ T cell immunotherapy.

In the current study, Vγ9+Vδ2+ γδ T cells did not expand after the second dose of ZOL+IL-2, which may indicate that these cells are refractory to a second phosphoantigen stimulation soon after the initial expansion *in vivo*. Monthly infusions of ZOL in pediatric leukemia patients has been shown to reduce circulating Vδ2+ γδ T cell numbers over time (35). Chen et al. showed in macaques that a second phosphoantigen dose, given 12 days later, induced a slight increase in the number of circulating Vγ9+Vδ2+ γδ T cells, although this increase was reduced in magnitude compared to the first dose (17). In contrast, when doses are separated by several months in anti-TB drug treated, Mtb-infected macaques, Vγ9+Vδ2+ γδ T cells re-expand significantly (29). These data suggest that shorter intervals of phosphoantigen expansion, as used here, may lead to Vγ9+Vδ2+ γδ T cells that are refractory to further expansion and could even lead to fewer of these cells.

While the transient induction of Vγ9+Vδ2+ γδ T cells in our study did not reduce TB disease, other studies using treatment strategies that promote a large, sustained presence of Vγ9+Vδ2+ γδ T cells in the lung during the early stages of Mtb infection may result in a better TB outcome. Shen and colleagues showed that immunization of macaques with a *Listeria monocytogenes* vaccine vector producing HMBPP, a potent stimulator for Vγ9+Vδ2+ γδ T cells, resulted in sustained expansion of these cells in blood and airways, and was associated with lower bacterial burden following Mtb infection (16). In this study, Vγ9+Vδ2+ γδ T cells remained significantly elevated 3 months after vaccination, which may reflect the higher potency of the HMBPP-mediated stimulation compared to ZOL + IL-2 stimulation (16). Qaqish and colleagues adoptively transferred Vγ9+Vδ2+ γδ T cells expanded *ex vivo* into SIV-naïve adult rhesus macaques at 3 and 18 days after Mtb infection, which resulted in lower Mtb burden (18). They noted early trafficking and retention of the adoptively transferred Vγ9+Vδ2+ γδ T cells to airway by 1 week after infection with Mtb (18). The reduction in Mtb burden observed in that study (18) may also be due to the large number of Vγ9+Vδ2+ γδ T cells transferred (∼10^8^ cells per infusion), which was far larger than the increases achieved here with *in vivo* ZOL+IL-2.

We observed alterations in the cytotoxic profiles of CD8αβ+ and CD8αα+ T cells, with higher frequencies of cells double- and triple-positive for granzyme B, perforin, and granulysin in the ZOL+IL-2-treated animals. Adjunctive ZOL+IL-2 co-administered with second-line anti-TB drugs has been shown to enhance cytotoxic CD8αβ+ effector T cells in airways of macaques infected with multidrug-resistant Mtb and led to lower bacterial loads than by antibiotics alone (29). However, in our study of SIV/Mtb coinfected juvenile macaques, while ZOL+IL-2 enhanced CD8αβ+ and CD8αα+ T cell cytotoxicity, it was not associated with a reduction in Mtb burden. In fact, thoracic lymph nodes tended to exhibit slightly more pathology in the ZOL+IL-2-treated group, indicative of greater lymph node involvement. There are several possible mechanisms by which ZOL+IL-2 may have enhanced CD8αβ+ and CD8αα+ T cell cytotoxic capacity in our study. The first is through direct cross-priming of CD8+ T cells. γδ T cells have been demonstrated to act as antigen presenting cells, thereby enhancing antigen-specific CD8+ T cell responses against various cancers and viral infections (36–39). The second is through indirect interactions. There is evidence that γδ T cells enhance dendritic cell priming of CD8+ T cells through cytokine secretion, resulting in enhanced antigen-specific CD8+ T cell responses (40). The third is a direct effect of the treatment itself. Expansion of Vγ9+Vδ2+ γδ T cells is dependent upon coadministration of ZOL and IL-2. (30). It is known that IL-2 itself can induce production of cytotoxic factors in CD8+ T cells (41, 42). Thus, the enhanced cytotoxic potential observed here may be a direct result of IL-2, although we did not have an IL-2-only group with which to test this possibility.

There are several limitations to the study presented here. We did not include animals treated with ZOL or IL-2 alone to identify effects attributable to each agent alone. In addition to influencing cytotoxic factors in CD8+ T cells, IL-2 alone can result in CD4+ T cell expansion and elevated plasma viremia (43, 44). However, we observed neither an increase in SIV viremia nor a higher frequency of circulating CD4+ T cells following ZOL+IL-2. Since evidence suggests that monotherapy with either ZOL or IL-2 does not expand Vγ9+Vδ2+ γδ T cells in humans or macaques (29, 45, 46), these extra single-agent control groups were not included. While potentially interesting, they would have doubled the size of our study which was not feasible. For similar reasons, we did not include an SIV-naïve group or an ART-treated, SIV-infected group, which may have generated a more vigorous γδ T cell response. To date, most NHP studies to assess the efficacy of γδ T cell immunotherapy have done so in the absence of preexisting SIV infection. We have shown previously that MCM chronically infected with SIV had defective adaptive immune responses to Mtb coinfection (47). Treating SIV-infected juvenile macaques with ART appeared to ameliorate these defects (26). Zhou and colleagues have also shown that SIV infection impairs mycobacterial-specific responses in Vδ2+ γδ T cells (48). Thus, the animals studied here may have had impaired immune responses to Mtb due to their preexisting SIV infection and the transient increase in Vγ9+Vδ2+ γδ T cells was unable to overcome that defect and restrict TB progression. Future studies that include ART treatment could better reveal the full potential of Vγ9+Vδ2+ γδ T cell restoration for controlling Mtb coinfection. Lastly, necropsies were performed 8 weeks after Mtb coinfection. This is the time at which the adaptive immune response to Mtb is maturing and the cells within granulomas begin to exert a sterilizing effect (49). Thus, the negligible difference in bacterial burden between groups that we observed here may be, at least in part, because adaptive immunity and mycobactericidal activity are just beginning to peak at 8 weeks. A longer follow-up period after Mtb coinfection may have revealed treatment related differences. However, in our previous studies of Mtb coinfection of SIV+ MCM, animals begin to develop extensive TB disease (*e.g.*, pneumonia and lung consolidations) after 8 weeks of coinfection (31) and this advanced pathology would have limited our ability to carefully immunophenotype T cells within individual granulomas. Here, we noted similar Vγ9+Vδ2+ γδ T cell frequencies in lung granulomas from ZOL+IL-2-treated and saline-treated animals. The necropsies 8 weeks after Mtb coinfection may have been too late to detect differences in migration or expansion of Vγ9+Vδ2+ γδ T cells in lung tissue, since circulating γδ T cells peaked within the first two weeks of Mtb coinfection. However, earlier necropsies would have reduced our ability to assess the efficacy of boosting Vγ9+Vδ2+ γδ T cells on mitigating TB disease.

Host-directed therapies focused on augmenting Vγ9+Vδ2+ γδ T cells are a promising strategy to mitigate Mtb burden and tissue damage, especially in children with HIV-associated TB. Our results suggest that ZOL+IL-2 induced a transient elevation of circulating Vγ9+Vδ2+ γδ T cells in SIV+ juvenile macaques when administered shortly after Mtb coinfection. However, this had little impact on TB disease or Mtb burden. Others have elicited more sustained increases of Vγ9+Vδ2+ γδ T cells over the course of Mtb infection in macaques and have shown an associated decrease in Mtb burden (16, 18). However, this has yet to be tested in the context of preexisting SIV. Despite the lack of efficacy observed here, augmenting Vγ9+Vδ2+ γδ T cells may provide measurable benefit as an adjunct to anti-TB chemotherapy in SIV+ animals based on data from SIV-naïve macaques (29). In fact, recent evidence suggests that adjunctively boosting Vγ9+Vδ2+ γδ T cells may reduce pathology and Mtb load in patients with multi-drug resistant TB (50). Boosting Vγ9+Vδ2+ γδ T cells with ZOL+IL-2 may also be beneficial when adaptive immunity is more fully developed, such as in latent TB infection. Further studies to elucidate the trafficking of Vγ9+Vδ2+ γδ T cells and their retention within various tissue compartments would also be useful for γδ T cell therapeutic strategies. Given the potential long-term health consequences of TB early in life, identifying a host-directed therapy that enhances Vγ9+Vδ2+ γδ T cell responses in combination with anti-TB chemotherapy may be a promising strategy, especially for children living with HIV who are at an elevated risk of TB.

## Methods

### Animals

Juvenile (∼1-2 years, equivalent to 4-8 years-old children) Mauritian cynomolgus macaques (*Macaca fascicularis*) were obtained from Bioculture US (Immokalee, FL) (Table S1). MHC haplotype was determined by MiSeq sequencing and animals with the presence of at least one copy of the M1 MHC haplotype were selected for this study (51), for consistency with our previous SIV/Mtb coinfection studies (26, 31, 47, 52, 53).

Animal protocols and procedures were approved by the University of Pittsburgh Institutional Animal Care and Use Committee (IACUC) which adheres to guidelines established in the Animal Welfare Act and the Guide for the Care and Use of Laboratory Animals, as well as the Weatherall Report (8th Edition). The University is fully accredited by AAALAC (accreditation number 000496), and its OLAW animal welfare assurance number is D16-00118. The IACUC reviewed and approved the study protocols 19014337 and 22010433, under Assurance Number A3187-01.

Animal welfare was monitored as described previously (26). In brief, all animals were checked at least twice daily to assess appetite, attitude, activity level, hydration status, etc. Following Mtb infection, the animals were monitored closely for clinical signs of TB (*e.g.*, weight loss, tachypnea, dyspnea, or coughing). Physical exams, including weights, were performed on a regular basis. Animals were sedated for all veterinary procedures (*e.g.*, blood draws) using ketamine or other approved drugs. Regular PET/CT imaging was conducted and has proven to be very useful for monitoring TB progression. Our experienced veterinary technicians monitored animals especially closely for any signs of pain or distress. If any were noted, appropriate supportive care (*e.g.*, dietary supplementation, rehydration) and treatments (analgesics) were given by trained staff. No animal on this study reached humane endpoint or required any intervention. At planned endpoint, each animal was heavily sedated with ketamine and humanely euthanized using sodium pentobarbital.

### SIV and Mtb infection

All animals were infected intravenously with 10,000 IU of SIVmac239M, a molecularly barcoded virus stock generated from clonal SIVmac239 (54). Six months later, animals were coinfected with low dose (17-18 CFU) of barcoded Mtb Erdman via bronchoscopic instillation as previously described (26) and followed for 8 weeks.

### Zoledronate + IL-2 treatment

Zoledronate (Reclast®; ZOL) was obtained from the University of Pittsburgh Medical Center Presbyterian Pharmacy and recombinant human IL-2 (IL-2) was obtained from Peprotech (Cat. No. 200-02). One group of SIV-infected juvenile animals (n = 5) received ZOL (0.2 mg/kg) by intravenous injection at day 3 and day 17 after Mtb coinfection. These animals then received daily IL-2 (0.2 mg/kg) subcutaneously for 5 days. The other group of SIV-infected juvenile animals (n = 5) served as controls which received saline alone at the ZOL timepoints.

### Clinical and microbiological monitoring

All animals were assessed twice daily for general health and monitored closely for clinical signs of TB (coughing, weight loss, tachypnea, dyspnea, etc.) following Mtb infection. Monthly gastric aspirates (GA) and bronchoalveolar lavage fluid (BALF) were cultured for Mtb growth. GA and BALF samples with culturable Mtb (+) or that were sterile (-) are indicated in Table S1. Blood was drawn at regular intervals as indicated to measure erythrocyte sedimentation rate (ESR) and to provide peripheral blood mononuclear cells (PBMC) and plasma for analysis.

### Viral loads

Plasma viremia was monitored serially by quantitative PCR as previous described (26). In brief, viral RNA was isolated using the Maxwell Viral Total Nucleic Acid Purification Kit (Promega, Madison, WI) and reversed transcribed using the TaqMan Fast Virus 1-Step qRT-PCR Kit (Invitrogen). DNA was quantified on a LightCycler 480 (Roche, Indianapolis, IN). Plasma viremia for both treatment groups is plotted in Figure S1.

### PET/CT imaging and analysis

Radiolabeled 2-deoxy-2-(^18^F)fluoro-D-glucose (FDG) PET/CT was performed just prior to Mtb infection and then monthly after Mtb infection. Imaging was performed using a MultiScan LFER-150 PET/CT scanner (Mediso Medical Imaging Systems, Budapest, Hungary) housed within our BSL3 facility as previously described (55, 56). Co-registered PET/CT images were analyzed using OsiriX MD software (version 12.5.2, Pixmeo, Geneva, Switzerland) to enumerate granulomas and to calculate the total FDG avidity of the lungs, exclusive of lymph nodes, which is a quantitative measure of total inflammation in the lungs (55, 57). Thoracic lymphadenopathy and extrapulmonary dissemination of Mtb to the spleen and/or liver were also assessed qualitatively on these scans.

### PBMC and BALF processing

PBMC were isolated from blood using Ficoll-Paque PLUS gradient separation (GE Healthcare Biosciences). Single-cell suspensions were cryopreserved in fetal bovine serum containing 10% DMSO in a liquid nitrogen freezer. BALF (2 x 10 mL washes of PBS) was pelleted and a 15 mL aliquot was cryopreserved. The cell pellets were resuspended into ELISpot media (RPMI 1640, 10% heat-inactivated human albumin, 1% L-glutamine, and 1% HEPES) and counted. BALF cells were then stained for flow cytometry.

### Necropsy

Necropsies were performed 8-9 weeks after Mtb infection as previously described (26). A final FDG PET/CT scan was performed within three days of necropsy to document disease progression and to guide the collection of individual granulomas (58). One saline-treated animal (166-21) was euthanized prior to necropsy due to complications during the 8-week (pre-necropsy) PET/CT scan. Thus, several analyses such as Mtb burden and immunophenotyping of tissues are unavailable for this animal. Animals for necropsy were heavily sedated with ketamine, maximally bled, and humanely euthanized using sodium pentobarbital (Beuthanasia, Schering-Plough, Kenilworth, NJ). Granulomas matched to the final PET/CT images were harvested along with thoracic and extrathoracic lymph nodes, lung tissue, as well as portions of liver and spleen. Quantitative gross pathology scores were calculated and reflect overall TB disease burden for each animal (58). Tissue samples were divided and a portion was fixed in 10% neutral buffered formalin (NBF) for histopathology; the remainder was homogenized to a single-cell suspension as described previously (58). Serial dilutions of these homogenates were plated onto 7H11 agar, incubated at 37°C, 5% CO_2_ for three weeks, and colonies were enumerated. Bacterial load in lungs, thoracic lymph nodes, liver, and spleen, as well as total thoracic CFU, were calculated as described previously (58). NBF-fixed tissue was embedded in paraffin, sectioned, and stained with hematoxylin and eosin for histopathologic examination.

### Flow cytometry

In general, cells collected from whole blood, BAL and necropsy were stained following a similar protocol: viability, Vδ2, surface and intracellular staining (for BAL and necropsy samples). A list of antibodies is provided in Table S2. For whole blood staining, samples underwent RBC lysis using 1X lysis buffer (BD Pharm Lyse ™ Lysing Buffer, Cat. No. 555899) in the dark for 8 minutes at room temperature (25°C). Cells were then stained for viability using a Live/Dead Fixable Aqua stain kit (Invitrogen, Cat. No. L34957) for 10 minutes at room temperature. Cells were washed, surface stained with freshly prepared, Zenon-labeled Vδ2 antibody in PE (Invitrogen, Cat No. Z25055) for 20 minutes in dark at room temperature, followed by surface staining (Table S2) for 20 minutes at 4°C. Cells were washed and then fixed for 10 minutes with 1% paraformaldehyde (PFA) in 1X PBS.

For BAL and necropsy tissues, cells were counted, reconstituted in ELISpot media (RPMI 1640 + 10% human albumin + 1% glycine + 1% HEPES buffer), aliquoted at 1×10^6^ cells/well in a 96-well plate and stimulated for 6 hours. For BAL, cells were either stimulated or not with 20 ng/mL of the γδ T cell stimulator (E)-4-Hydroxy-3-methyl-but-2enyl pyrophosphate (HMBPP; Sigma-Aldrich, Cat. No. 95098-1MG). CD107a-BV421 (Biolegend), CD154/CD40L-PE-Dazzle594 (Biolegend), brefeldin A (1 μg/mL; eBioscience, Cat. No. 00-4506-51) and monensin (0.5 μM; Biolegend, Cat. No. 420701) were added as well. After 6 hours, cells were washed and stained for viability (Live/Dead Aqua; Invitrogen, Cat. No. L34957). Vδ2 γδ T cells and surface (Table S2) were stained the same as whole blood and samples were fixed in 1% PFA. Cells were then permeabilized for 10 minutes using a BD Cytofix/Cytoperm™ kit (BD, Cat. No. 554714), stained with intracellular staining cocktail for 20 minutes in the dark at room temperature, washed, and analyzed. For necropsy tissues, cells isolated from granulomas, spleen, and PBMC were stimulated for 6 hours. Spleen and PBMC were stimulated with PDBU and ionomycin. In brief, stimulators were added and incubated for 1 hour, then brefeldin A (1 μg/mL) was added for the remainder of the stimulation time. Cells were stained with a Live/Dead Blue Fixable dye (Invitrogen, Cat. No. L23105) for 10 minutes at room temperature. Cells were washed, incubated with Vδ2 antibody at room temperature for 20 minutes, and then incubated with AlexaFluor 647-labeled goat anti-mouse IgG1 (1:500; Invitrogen, Cat. No. A21240) for 20 minutes. Cells were stained with surface antibody cocktail (Table S2) for 20 minutes at 4°C, then fixed in 1% PFA and permeabilized with BD Cytofix/Cytoperm™ (BD; Cat No. 554714). Cells were then stained intracellularly for 20 minutes at room temperature, washed, and analyzed.

Flow cytometry was performed using a Cytek Aurora (BD). FCS files were analyzed using FlowJo software for Macintosh (version 10.1). Gating strategies for whole blood, BAL and necropsy data are shown in Figure S3, S4, and S6, respectively. For most samples, we acquired 50,000 events in the lymphocyte gate. When this was not possible (*i.e.*, for some small granulomas), we applied a cutoff threshold of CD3 events >100. Samples below that threshold were excluded from further analysis. For 159-21 and 162-21 necropsy data, Vδ1 and granulysin were excluded from the analysis due to the addition of granulysin-PE-Cy7 to the antibody cocktail during staining, impeding the ability to analyze Vδ1 γδ T cells, which also used an antibody conjugated to PE-Cy7. These samples were gated similar to as shown in Figure S6.

### Statistics

For comparing longitudinal plasma viremia data, a linear mixed model with subject as a random variable were used to test treatment groups over time. Fixed effect tests were used to assess whether there were differences among treatment groups or among time points. Time points after Mtb were then compared within each treatment by Dunnett’s multiple comparisons test relative to the ‘Pre Mtb’ time point. For all other data, the Shapiro-Wilk normality test was used to check for normal distribution of data. Unpaired normally distributed data were analyzed using t tests, while unpaired non-normally distributed data were analyzed with the Mann-Whitney U test. All statistical tests were performed in Prism (version 9.0.0; GraphPad). All tests were two-sided, and statistical significance was designated at a *P* value of < 0.05.

## Acknowledgements

We thank the incredible veterinary and laboratory staffs of the TB Research Group at the University of Pittsburgh for their contribution to this study. We are grateful to Dr. Edwin Klein, DVM, for his expert review of the histopathology slides. E.C.L was supported by National Institute of Health (NIH) K01 award OD033539. This work was supported by NIH R01 AI142662 awarded to L.C.N., S.L.O., and C.A.S. D.I.G. was supported by a Senior Research Fellowship (1117766) and subsequently by an NHMRC Investigator Grant (2008913). NIH award (UC7AI180311) from the National Institute of Allergy and Infectious Diseases (NIAID) supported the operations of the University of Pittsburgh Regional Biocontainment Laboratory (RBL) within the Center for Vaccine Research (CVR). **Disclaimer:** The opinions expressed in this article are the authors’ own and do not necessarily reflect the view of the National Institutes of Health, the Department of Health and Human Services, or the United States government.

---

**Figure S1. Plasma viremia for saline and ZOL+IL-2.** Plasma viral load (viral copy equivalents/mL) were determined by qRT-PCR. Each point indicates an individual animal. Triangles along x-axis indicate when treatment was administered (days 3 and day 17 post Mtb). Horizontal dashed line represents the limit of detection.

**Figure S2. Frequencies of CD4+ & CD8+ T cell subsets in blood and BAL after Mtb coinfection and ZOL+IL-2/saline treatment.** Lines indicate mean frequencies and error bars indicate standard deviation of treatment groups. Grey triangles along x-axis indicate when treatment was administered (days 3 and day 17 post Mtb).

**Figure S3. Gating schematic for whole blood panel.** Whole blood samples were stained for flow cytometric analysis as indicated in the Methods using antibodies listed in Table S2 to determine the T cell subsets over SIV infection, Mtb coinfection, and ZOL+IL-2/saline treatment. Total whole blood samples were gated for live cells followed by lymphocytes. B cells and small monocytes/macrophages were removed (CD20 & CD163 vs. SSC-H) followed by singlets and then T cells (CD3+). From the T cell gate, Vδ1 γδ T cells were gated. Vδ1-T cells were characterized by Vγ9 and Vδ2 expression. Vγ9-Vδ2-T cells were further categorized for CD4+ or CD8+ T cells.

**Figure S4. Gating schematic for BALF panel.** Fresh BALF cells were stimulated for 6 hours +/- HMBPP, a γδ T cell stimulator. After 6 hours, cells were stained for flow cytometric analysis as indicated in the Methods using antibodies listed in Table S2 to determine γδ, CD4+ and CD8+ T cell subsets in airways over SIV infection, Mtb coinfection, and ZOL+IL-2 or saline treatment. A) Total BAL cells were gated for live cells followed by leukocytes (CD45 vs. SSC-B-H). Then, leukocytes were sub-gated by lymphocytes and myelocytes. From the lymphocyte gate, B cells and macrophages were removed (CD20 & CD163 vs. SSC-H) followed by singlets and then T cells (CD3+). From the T cell gate, Vδ1 γδ T cells were gated. Vδ1-T cells were characterized by Vγ9 and Vδ2. Vγ9-Vδ2- were further categorized by CD4+ or CD8+ T cells. B) All T cell subsets were further characterized for phenotype and function. The CD8 T cell population was used here as a representative population for gating. Memory was characterized: Naïve T cells (Tn, CD95-CD28+); Transitional memory T cells (Ttm, CD95+CD28+); and Effector memory T cells (Tem, CD95+CD28-). Cytokine responses were gated on IFNγ or TNF versus CD69. CD107a, CD154, HLA-DR, PD-1, ICOS, and Granzyme B (GrzB) were also characterized.

**Figure S5. *De novo* cytokine production of CD8 T cell subsets in granulomas.** Granulomas were incubated for 6 hours for *de novo* cytokine production. Outlined symbols indicate median per animal and unlined symbols indicate individual samples. Bars indicate group medians. Two-tailed, Mann Whitney U tests were performed to determine significance. P-values are shown. A) IFNγ production in CD8ɑβ+ T cells. B) TNF production in CD8ɑβ+ T cells. C) IL-2 production in CD8ɑβ+ T cells. D) IL-17 production in CD8ɑβ+ T cells. E) IFNγ production in CD8ɑɑ+ T cells. F) TNF production in CD8ɑɑ+ T cells. G) IL-2 production in CD8ɑɑ+ T cells. H) IL-17 production in CD8ɑɑ+ T cells.

**Figure S6. Gating schematic for necropsy panel.** Cells isolated from granulomas were incubated for 6 hours after which cells were stained for flow cytometric analysis as indicated in the Methods using antibodies listed in Table S2 to determine γδ, CD4+ and CD8+ T cell subsets as well as phenotype and function. A) Total cells were gated for singlets, then live cells, then lymphocytes, followed by B cells (CD20+CD3-) and T cells (CD20-CD3+). From the T cell gate, Vδ1 γδ T cells were gated. Vδ1-T cells were characterized by Vγ9 and Vδ2. Vγ9-Vδ2- were categorized by CD4+ or CD8ɑ+ T cells. CD8ɑ+ T cells were gated further on CD8β+ (CD8ɑβ) or CD8β- (CD8ɑɑ). B) All T cell subsets were characterized for phenotype and function. The CD8 T cell population was used here as a representative population for gating. Memory was characterized: Naïve T cells (Tn, CD95-CD28+); Transitional memory T cells (Ttm, CD95+CD28+); and Effector memory T cells (Tem, CD95+CD28-). Cytokine responses were gated on IFNγ, TNF, IL-2, and IL-17 versus CD69. Cytotoxic factors were gated: Granzyme B (GrzB), Granulysin (Glyn), and Perforin (Perf).

**Figure S7. ZOL+IL-2 does not reduce lung inflammation during Mtb coinfection.** Total lung FDG activity relative to weeks after Mtb coinfection. Lines indicate mean FDG activity and error bars indicate standard deviation of treatment groups.

**Table S1.**
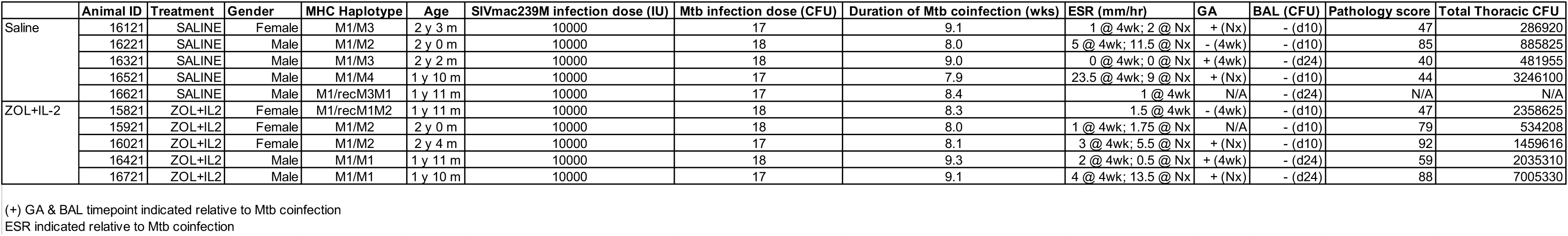
Summary of juveniles and outcome measures following Mtb coinfection. For erythrocyte sedimentation rate (ESR), gastric aspirate (GA), and bronchoalveolar lavage (BAL), time point indicated relative to Mtb coinfection.

**Table S2.**
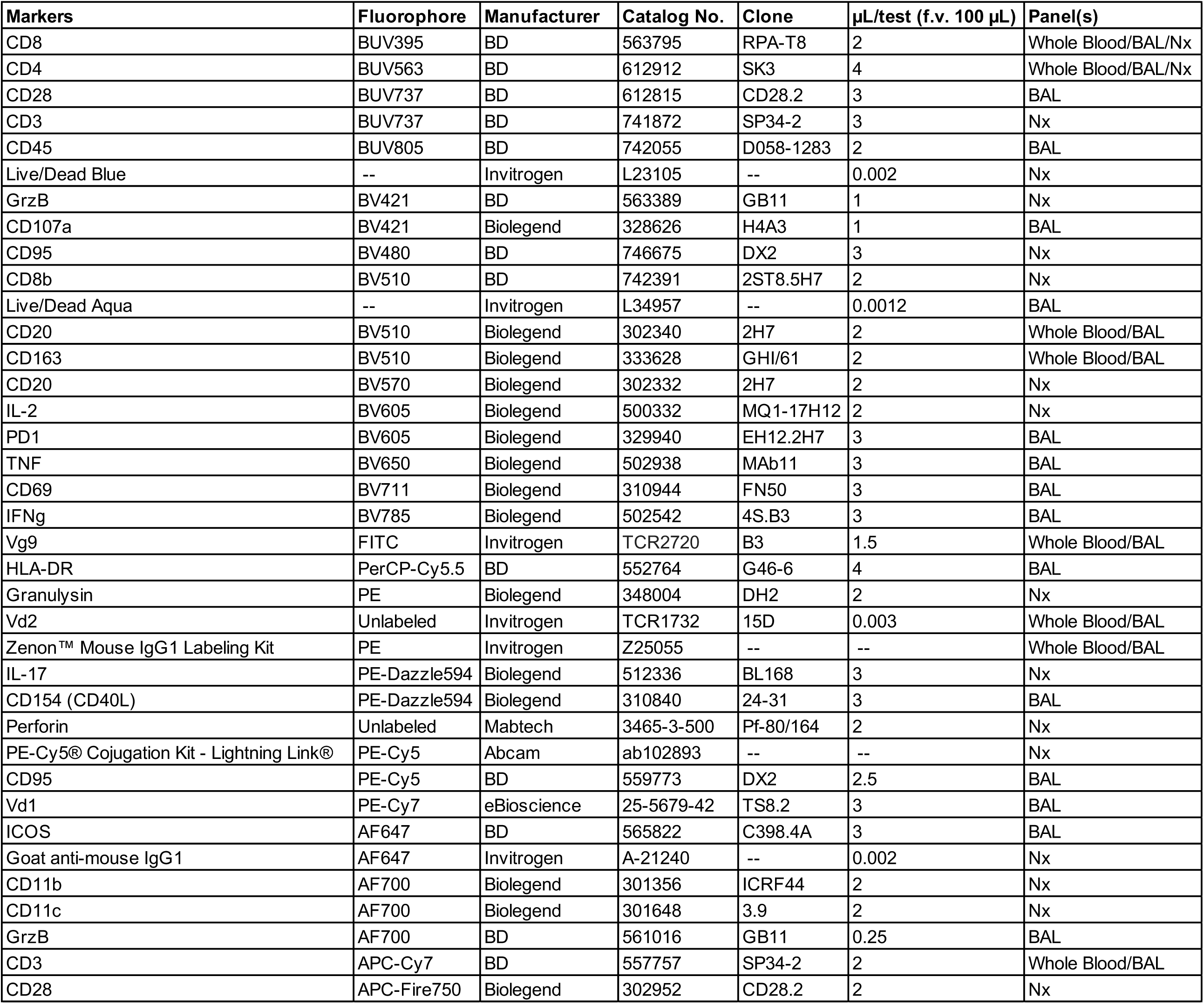
Antibody list by flow panel. All flow staining was performed in final volume (f.v.) of 100 µL.

